# Unified machine learning framework uncovers overlooked bias control in immunogenic neoantigen identification

**DOI:** 10.1101/2024.02.07.579420

**Authors:** Ziting Zhang, Wenxu Wu, Lei Wei, Hai Qi, Xiaowo Wang

## Abstract

Neoantigens play a crucial role in tumor immune process and precisely identifying them can greatly contribute to tumor immunotherapy design. There are three main steps in the neoantigen immune process, i.e., binding with MHCs, extracellular presentation, and immunogenicity induction. Various computational methods have been developed, but the overall accuracy of neoantigen identification remains relatively low. Here, we established a unified transformer-based framework ImmuBPI that comprised three tasks. Cross-task model interpretation discovered a counterfactual pattern learned by the immunogenicity prediction model overlooked in previous studies. We demonstrated that this model bias arose from training data imbalance and the neglect of bias control would not only mislead the models but also hinder existing benchmarks from fair evaluation. We further designed a mutual information-based debiasing strategy that showed preliminary potential in alleviating the bias. We believe these observations will provide insightful perspectives for future neoantigen prediction by highlighting the necessity of bias control.

## 1 Introduction

Tumor immunotherapy is a promising, rapidly growing generation of cancer treatment that aims to kill cancer cells by leveraging one’s own immune system [1]. A range of immunotherapies have already been approved, bringing significant clinical benefits to cancer patients [2, 3]. To achieve precise killing effects, the immune system relies on specific markers called neoantigens. They are a subset of abnormal peptides originating from genetic mutations within tumor cells. Accurately identifying neoantigens from vast tumor mutations can significantly contribute to immunotherapy development [4, 5]. There are three main steps in the neoantigen immune process, i.e., binding with Major Histocompatibility Complexes (MHCs), which is also called Human Leukocyte Antigen (HLA) in humans, extracellular presentation, and induction of immunogenicity by forming interactions with the T cell receptor (TCR) [6].

In recent years, a plethora of machine learning methods have been developed to predict the above three processes related to neoantigens, i.e., binding [7, 8, 9, 10, 11, 12, 13, 14], presentation [15, 16, 17, 18, 19, 20] and immunogenicity [21, 22, 23, 24, 25, 26, 27, 28, 29], each of which contributes to neoantigen identification from different perspectives. However, despite the progress made in both scientific research and clinical therapeutics [30, 31, 32, 33, 34], the overall accuracy of neoantigen identification remains far from satisfactory [35, 36, 37]. Only 6% of the predicted epitopes from the global consortium TESLA were positive when tested with immunogenicity [36].

One limitation in this field is the lack of a systematic view and understanding of the complicated, multi-step processes involved in neoantigen identification. Contrary to the intensive focus on improving model performance for individual tasks, few studies have explored relationships among models trained for diverse immunological tasks (Fig. 1A). There is still ongoing controversy regarding the optimal way to integrate models for different tasks in real-world neoantigen identification[36, 37]. Another challenge arises from substantial data bias, caused by inevitable batch effect and high experimental cost with immunological assays [5, 38] (Fig. 1B). In biological scenarios, biased data distribution is a common phenomenon[39, 40, 41, 42, 43, 44]. It has been reported that this can lead data-driven models to find shortcuts thus making spurious decisions when generalized to test sets [45, 46, 47]. Nevertheless, to the best of our knowledge, there have been no attempts to examine the potential effects of data bias in the neoantigen prediction field.

**Fig. 1.**
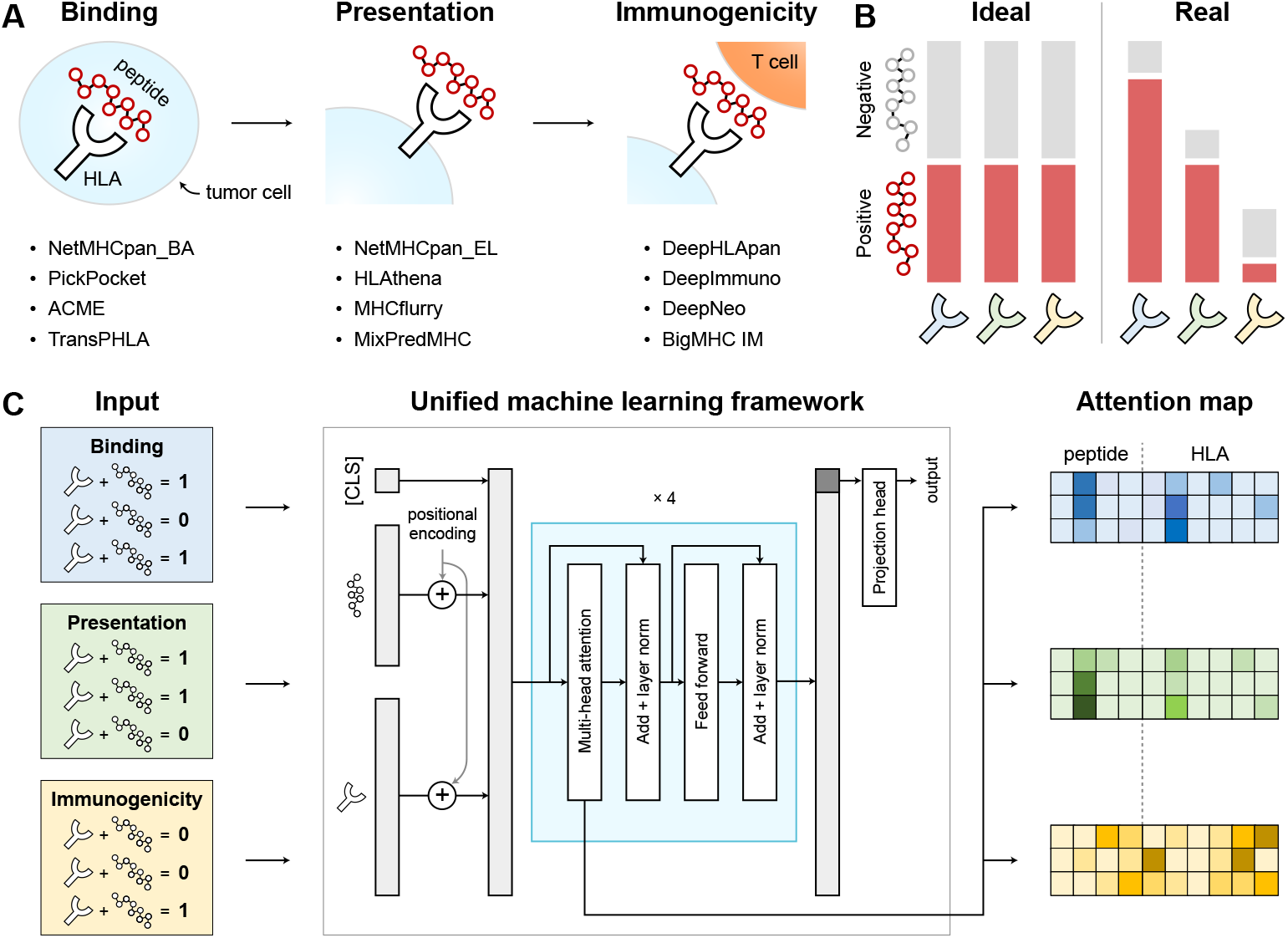
(A) Various models have been applied to model three immunological tasks involved in neoantigen predictions, yet limited studies have explored relationships among models trained for diverse immunological tasks. (B) An ideal distribution (left) should achieve a balance in both the number of peptides related to different HLA alleles and the ratio of positive peptides to negative peptides given a specific HLA allele. Dataset distributions in real-world scenarios (right). (C) The overview of ImmuBPI as a unified machine learning framework. After training for different tasks, the attention map of each model was displayed and compared to verify the model’s major determinants.

In this paper, we established a unified machine learning framework (Fig. 1C), named ImmuBPI (**Immu**ne predictors for **B**inding, **P**resentation, and **I**mmunogenicity), to model three key steps of the neoantigen immune response process. ImmuBPI demonstrated superior predictive power with attention mechanisms as a powerful model indicator. Through interpretability analysis across different tasks, we have identified a significant bias in the immunogenicity dataset overlooked in previous studies. We illustrated that this bias would lead data-driven models to learn skewed discriminatory boundaries in a model-agnostic way. To address the challenge, we developed a mutual information-based debiasing strategy that enabled the model to perform sensitive immunogenicity predictions on mutation variants, a task where current state-of-the-art methods fell short. After debiasing, clustering immunogenic peptides with immunogenicity-encoded representations uncovers unique preferences for biophysical properties, such as hydrophobicity and polarity. These observations serve as a key complement to the past understanding that accurately predicting neoantigen is constrained by limited data, highlighting that the neglect of bias control may exert a significant defect on the model’s generalization performance. We expect this study will provide novel and insightful perspectives for developing neoantigen prediction methods and benefit future neoantigen-mediated immunotherapy designs.

## 2 Results

### 2.1 ImmuBPI learns three immunological tasks with a unified attention-based model architecture

In this work, we focused on three pivotal immunological processes: binding, presentation, and immunogenicity. The occurrence of each of the three events can be formulated as a binary classification problem based on the provided HLA and peptide sequence. The primary motivation was to make our analysis as universal as possible with popular and standard model architecture and to leverage the attention mechanism as an analysis method. We chose a transformer encoder-based model backbone to build the ImmuBPI framework [48] (Fig. 1C). The three models trained for binding prediction, presentation prediction, and immunogenicity prediction were correspondingly named ImmuBPI_B, ImmuBPI_P, and ImmuBPI_I.

To achieve a high classification performance, we chose an interaction-based model rather than a representation-based model [49], which was adopted by the current state-of-the-art model TransPHLA[14] in the binding prediction task. Besides, we employed an attention-pooling mechanism similar to that of Ref [50]. This involved a virtual and learnable token [CLS] (Fig. 1C) along with its associated embedding. The contextual embedding of this token was directly adopted as the representation for the entire sequence pair, serving as the input to the projection head for classification. These above modules provided an adaptive and multi-layer fusion of the peptide and HLA representation.

In addition to better benchmark performance, utilizing a transformer encoder-based model yielded the attention map thus offering the advantage of good interpretability. By extracting the row corresponding to the [CLS] token as a query from the attention map, we gained insights into how other parts of the input contributed to the embedding implicitly learned from training.

### 2.2 ImmuBPI demonstrates superior performance in binding and presentation prediction via learning anchor residues-HLA interactions

To systematically evaluate ImmuBPI’s effectiveness in binding prediction, we adopted the same setting and two benchmarks from the latest state-of-the-art methods [14]. Overall, 9 models [8, 11, 14, 16, 51, 52, 53, 54] were engaged in the benchmark, including the current state-of-the-art method TransPHLA. Because not all methods can be applied universally to peptides of all lengths and HLA alleles, performance was compared only when the respective model could successfully predict all the data in the benchmark to ensure a fair comparison. All comparative models’ performances were directly retrieved from Ref [14]. ImmuBPI_B achieved the best performance on both benchmarks on various metrics, with compatibility for all data pairs (Fig. 2A).

**Fig. 2.**
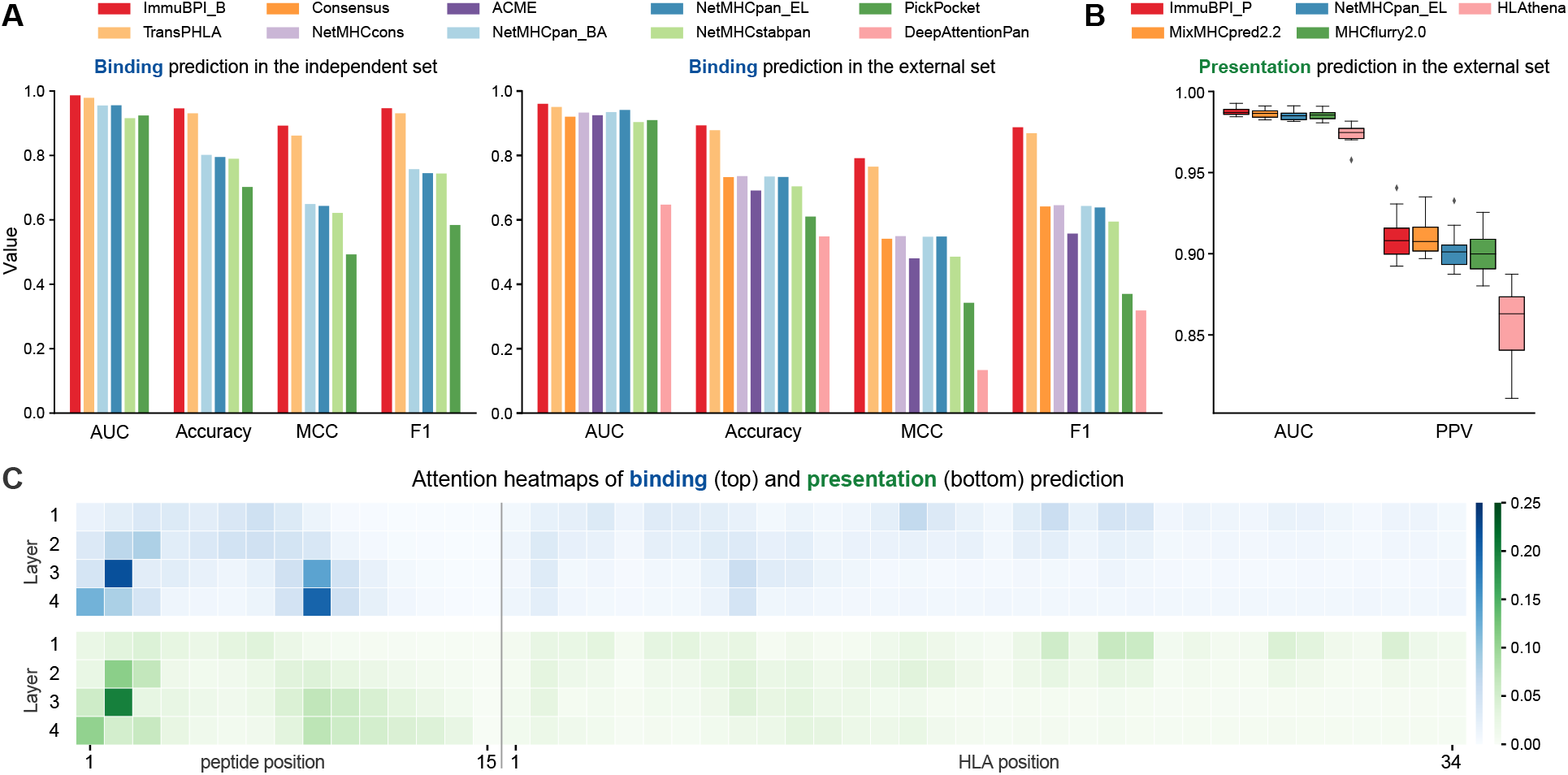
(A) Performance of different models in two test sets for the binding prediction task. Performance was compared only when the respective model could successfully predict all the data in the benchmark to ensure a fair comparison. (B) Performance of different models in the test set for the presentation prediction task. The value was plotted using boxplots as the test set contained ten samples. (C) Visualization of attention weights in heatmaps for binding and presentation prediction. The heatmap showed the contribution of each position on the peptide and HLA sequence to the [CLS] token. The four transformer encoder layers were arranged from top to bottom. Attention weights were averaged between different attention heads and ensemble models.

For presentation prediction, we utilized the same setting and independent benchmarks from the latest state-of-the-art methods [20]. The external test set consisted of HLA-I peptidomics from ten meningioma samples [55]. We added an extra post-processing step by assigning the highest score of all the peptide-HLA pairs possible in the sample to each peptide as the final score [20]. Overall, 4 models [16, 18, 19, 20] were tested in the benchmark, including the current IEDB [56] recommended method NetMHCpan EL and the state-of-the-art method MixMHCpred2.2. As shown in Fig. 2B, ImmuBPI_P had the highest AUC value and a comparative PPV value with MixMHCpred2.2, demonstrating the powerful predictive performance of our transformer-based framework.

To further investigate the model’s determinants, we performed the inference on the validation set and plotted the heatmap that visualized the attention weight of each amino acid from peptide and HLA respectively to the [CLS] token (Fig. 2C, Fig. S1). Both attention maps strongly highlight the first two and the last positions of the peptide, with an emphasis on the 9th position because 9mer constitutes the most common length. These positions are referred to as anchor positions, which are supported by previous studies [57, 58]. For allele-wise analysis, we selected 10 HLA types (4 HLA-A, 4 HLA-B, and 2 HLA-C) prevalent in the global population [59]. The sum of attention weights on 9-mer epitopes with positive binding or presentation label given a HLA allele were visualized in heatmaps [14] (Fig. S2). ImmuBPI can successfully reveal binding and presentation patterns for specific HLA alleles, which can be rather diverse for different HLA families. For example, while A2 family has a preference for peptides with a large, aliphatic side chain at the second position, B44 family favors the glutamic acids with a negative charge at the same position. These results align with previous findings [60, 61, 14] and prove the effectiveness of the attention mechanism for model interpretation.

To sum up, both binding and presentation are two important prerequisites for immunogenicity, which are to a large extent defined by interactions between anchor residues and HLA alleles. Binding predictions focus on the biochemical interaction between peptides and HLA molecules, whereas presentation predictions consider additional factors such as stability [51]. Due to abundant training data, the underlying interaction pattern is now relatively well-captured by ImmuBPI and other state-of-the-art models with high predictive performance.

### 2.3 Abnormal attention map for immunogenicity prediction task suggests hidden bias captured by current machine learning models

The ability to trigger immune responses by interacting with TCRs is referred to as immunogenicity. Contrary to the binding and presentation where anchor residues of the peptide are the main drivers, the non-anchor part, usually exposed at the interface, provides crucial signals for immunogenicity [62, 63]. As a result, HLA molecules and/or anchor residues are often excluded when considering T cell interactions in computational modeling frameworks [20, 25, 62, 64, 65]. With the expectation that models can automatically learn feature importance to generate informative representations, machine learning methods have recently been applied to immunogenicity predictions [23, 24, 27, 28] using both HLA allele and full-length peptide as input. However, to what extent the expectation holds in the context of immunogenicity prediction is largely unexplored.

In this section, we first conducted an in-depth characterization of the data distribution within the immunogenicity dataset. We collected training datasets from four recent machine learning-based immunogenicity prediction research [23, 24, 27, 28]. Significant imbalance exists from both inter-HLA and intra-HLA perspectives. Few HLAs with high frequency can make up a large portion of the training set (Fig. 3A). Another serious concern arises from the substantial intra-HLA imbalance, with over 50% of HLA alleles exhibiting a tenfold or greater imbalance between positive and negative peptides in all four datasets. This makes the common strategy designed for dataset imbalance infeasible (Fig. 3B).

**Fig. 3.**
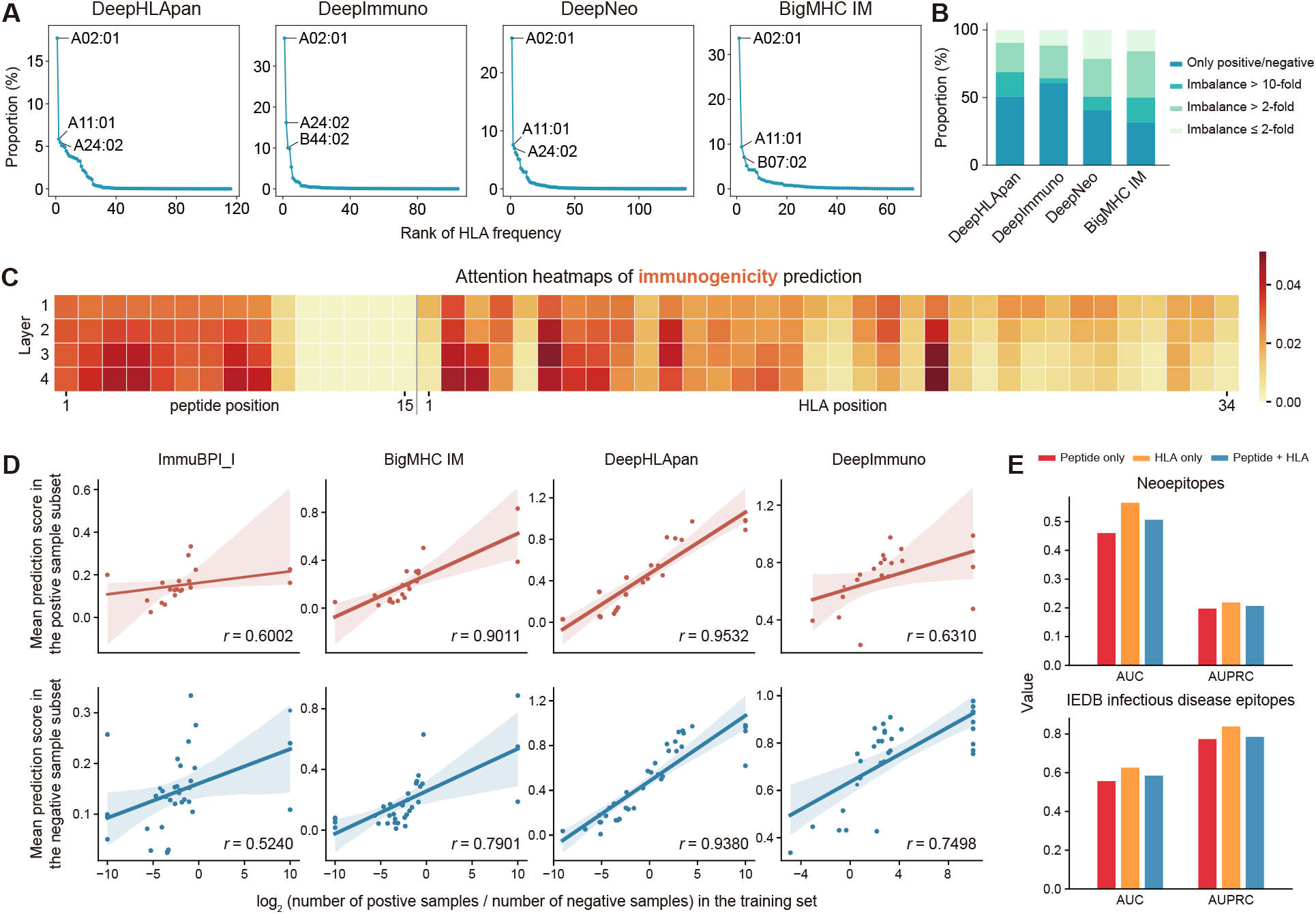
(A) Visualization of HLA frequency in different training datasets. X value represented the rank of HLA frequency. Y value represented the proportion of data points covered by the corresponding HLA allele. The three most frequent HLAs in each dataset are annotated in the figure. (B) Visualization of different training datasets in stacked bar charts. For each HLA allele in the training set, we calculated the ratio between the number of positive and negative peptides. HLA alleles were divided into four groups based on the imbalance degree. (C) Visualization of attention weights in heatmaps for immunogenicity prediction. (D) Correlation between model prediction scores in the neoepitopes benchmark and HLA imbalance degree in the training set. X value was calculated as the log2 ratio between positive and negative peptides corresponding to a given HLA allele in the training set. X value of HLA with only positive peptides was set at 10 while HLA with only negative peptides was set at −10. Y value was calculated as mean prediction scores across all peptide-HLA pairs related to this HLA. HLA with only one sample in the subset was excluded. (E) Performance of models trained with different inputs in two benchmarks.

Then, we moved on to investigate the impact of the observed imbalance on the models. Based on ImmuBPI_I, the attention map for immunogenicity prediction was plotted. HLA demonstrated abnormally high attention weights compared with binding and presentation results (Fig. 3C, Fig. 2C), contradicting the aforementioned biological facts. As the ratio between the number of positive and negative peptides given a specific HLA allele constitutes a highly variable feature for different HLA alleles, one conceivable hypothesis is that the model may learn shortcuts to make decisions using this feature.

To verify the assumption, we adopted two benchmarks from Ref [28] and each was further divided into two subsets based on the label. Within each subset, we investigated the correlation between positive-negative ratios for each HLA in the training set and the mean prediction score in the test set across all peptide-HLA pairs related to this HLA. Additionally, we performed the same analysis with BigMHC IM [28], DeepHLApan [23], and DeepImmuno [24] models with their respective training datasets to test whether our finding is model-agnostic. A significantly high Spearman correlation coefficient was consistently observed with all models in both benchmarks (Fig. 3D, Fig. S3A).

We also performed a mutation-based model-agnostic analysis [27] for four different machine learning models including ImmuBPI, DeepHLAPan, BigMHC IM, and DeepImmuno (Fig. S4). In binding and presentation tasks, the HLA and anchor positions showed the largest differences, which matched with the biological mechanism. Nevertheless, in all four machine learning models trained on the immunogenicity datasets, the HLA allele exhibited the largest variances compared with the peptide residues, which in fact contained the key signal for immune recognition. This experiment also supported our observations regarding the model bias.

These observations raised concerns about the capabilities of current immunogenicity benchmarks to fairly evaluate different models’ performance associated with the learning of immunogenicity. As current immunogenicity benchmarks are also composed of multiple retrospective studies, a similar distribution of intra-HLA imbalance between the training set and the benchmark may lead to an inflated performance and vice versa. We performed an ablation study and trained the ImmuBPI_I model using 3 different inputs: (1) only use peptides; (2) only use HLA alleles; (3) use both peptides and HLA alleles. Strikingly, the model achieved the best performance on both benchmarks when only the HLA allele was used. The performance of HLA-only model on IEDB benchmark had greatly surpassed (AUROC:+3.18% ∼ +15.95%, AUPRC:+5.13% ∼ +14.01%) all 15 models evaluated in Ref [28] (Fig. 3E).

In summary, we found that unbalanced data distribution broadly exists in various existing training datasets for immunogenicity prediction. This causes the decision boundaries of machine learning models to a considerable amount impacted by intra-HLA imbalance, which is a biologically irrelevant feature that varies among different datasets (Fig. S3B). Unfortunately, model performance evaluated by existing benchmarks may fail to precisely inflect its true ability to recognize immunogenicity. The real-world scenarios for immunotherapy design often require prioritizing the immunogenic peptides given a patient’s HLA typing [5], a setting that exhibits a substantial data distribution gap compared with existing datasets. We emphasize that the observation of both the model bias and benchmark design deserves more attention.

### 2.4 Mutual information-based debiasing strategy disrupts shortcut learning and enables sensitive immunogenicity prediction on mutation variants

The ideal approach to reduce the bias would be to have a large-scale, evenly distributed training dataset with little batch effect. Unfortunately, this solution is far from feasible due to high experimental cost, diverse experimental settings, unattainable uniform sampling strategy, and others [45]. To address the challenge, we used the mutual information-based debiasing strategy and showed its effectiveness through two case studies.

The fundamental premise of our debiasing strategy is to mask features that exhibit relatively high MI with the ground truth label but lack a strong biologically meaningful causal relationship with it. To first verify MI score aligned with the data imbalance that contributed to the bias, we proposed that the MI score between HLA and immunogenicity may serve as a quantitative metric for characterizing dataset imbalance. In datasets where HLA alleles with a substantial disparity between positive and negative samples predominate, the conditional entropy of the label given each HLA allele is consequently low, resulting in a high overall MI score. Calculations of MI scores in four datasets were consistent with previous findings shown in a qualitative way using stacked bar charts (Fig. 4A, Fig. 3B). Considering the non-dominant rule HLA plays in interaction with immune cells [62, 25, 20, 64, 65], it should be masked before input to prevent the model from learning the biased solution.

**Fig. 4.**
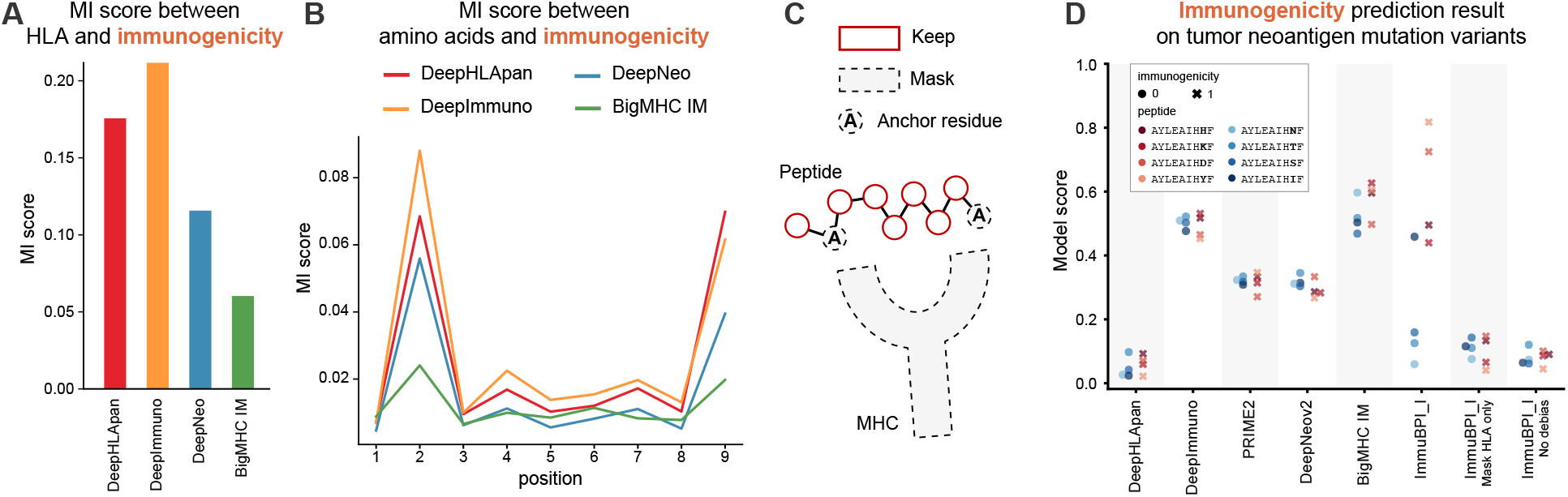
(A) MI score between HLA alleles and immunogenicity labels in four training sets. (B) MI score between amino acids on different positions of the peptide and immunogenicity labels in four training sets. (C) Debiasing strategy illustration. HLA alleles, the second and last anchor positions were masked before input. (D) Prediction results of different models on SNV mutations of a colorectal cancer neoantigen.

Intriguingly, though distribution related to HLA is the root of data bias, its removal alone does not offer a complete solution to the issue. Similar to the process above, we plotted the MI score between amino acids on different positions of the 9mer peptide and the immunogenicity label [23, 24, 27, 28] (Fig. 4B). The second and the last positions showed the highest MI scores on all four datasets despite their secondary role involved in TCR interactions [62, 63]. Based on previous analysis, peptides bound to different HLAs have conservative yet distinct patterns. As a result, the anchor position would to some degree leak the HLA information. In cases where strong interdependencies and correlations exist among features, determining the independent contributions of each feature by interpreting models can be challenging. We emphasize that the MI score is an informative complementary approach to model-based interpretability methods.

Finally, we added masks to both HLA alleles and two aforementioned anchor positions of the peptide and then trained the prediction model (Fig. 4C). We adopted a case study of non-synonymous single-nucleotide variants (SNVs) from a tumor neoantigen [66]. SNVs are a well-characterized category of neoantigens that has the potential to trigger effective immune responses [67, 68]. It can also lead to immunogenicity loss, referred to as immune escape [69]. Accurately identifying mutation variants with dramatic changes in immunogenicity is critical to immunology research and therapeutics design.

We compared ImmuBPI_I with five recently published models trained for immunogenicity prediction tasks [20, 23, 24, 27, 28]. Only ImmuBPI_I after debiasing demonstrated significant prediction variances and improved performance on mutation variants (Fig. 4D, Table 1). Interestingly, at least 6 out of 8 test data points actually appeared in the training set of PRIME2 and BigMHC IM [20, 28], yet the two models failed to make distinctive predictions, suggesting data bias may have a profound impact on the model’s decision boundaries. To make the results more generalizable, we also adopted a SARS-CoV-2 case from Ref [70] which shared a similar immune mechanism with tumor neoantigen, differing in the source of peptides. A similar trend can be found as our debiased model demonstrated higher sensitivity of prediction on mutations that changed immunogenicity (Fig. S5). Nevertheless, even after debiasing, the model failed to make precise predictions on 2 groups of the SARS-COV-2 case, suggesting the difficulty of the identification of immunogenicity.

**Table 1.**
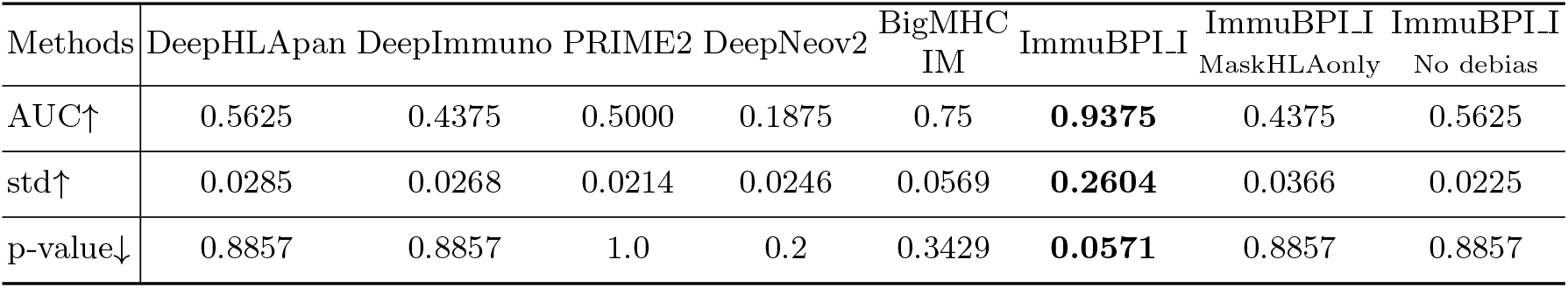
Performance of different models on SNV mutations of a colorectal cancer neoantigen, including AUC score on all mutants prediction, standard deviation (std) of all mutants prediction, and p-value of Mann–Whitney U Test on prediction scores of positive mutants versus prediction scores of negative mutants.

Admittedly, there would be information loss to some degree considering HLAs are also involved in TCR interactions. Employing a hard-masking strategy would make the model fail to distinguish different peptide-HLA pairs when only the HLA is different. Still, these results prove the necessity of bias control and that our debiasing strategy’s effectiveness in helping the model learn informative signals that decide peptide immunogenicity.

### 2.5. Clustering immunogenic peptides with immunogenicity-encoded representations uncovers unique preferences for biophysical properties

The improved performance on the immunogenicity test suggested that ImmuBPI_I can effectively capture the immunogenicity semantics associated with immunogenic epitopes. We extracted embeddings of the last projection layer of positive 9mer epitope populations in the training set. To identify biologically relevant peptide clusters, we used PCA to reduce dimensions to 20 and applied the K-means algorithm to perform clustering for ImmuBPI_I both with and without debiasing. We set K=7 according to the elbow plot and used PCA to further embed the sequence in 3-dimension to visualize the latent space (Fig. 5A). Without debiasing, the embedding space appeared to be less smooth and showed high consistency with the distribution of HLA typing. This again confirmed the previous conclusion that HLA was one of the major determinants of the model.

**Fig. 5.**
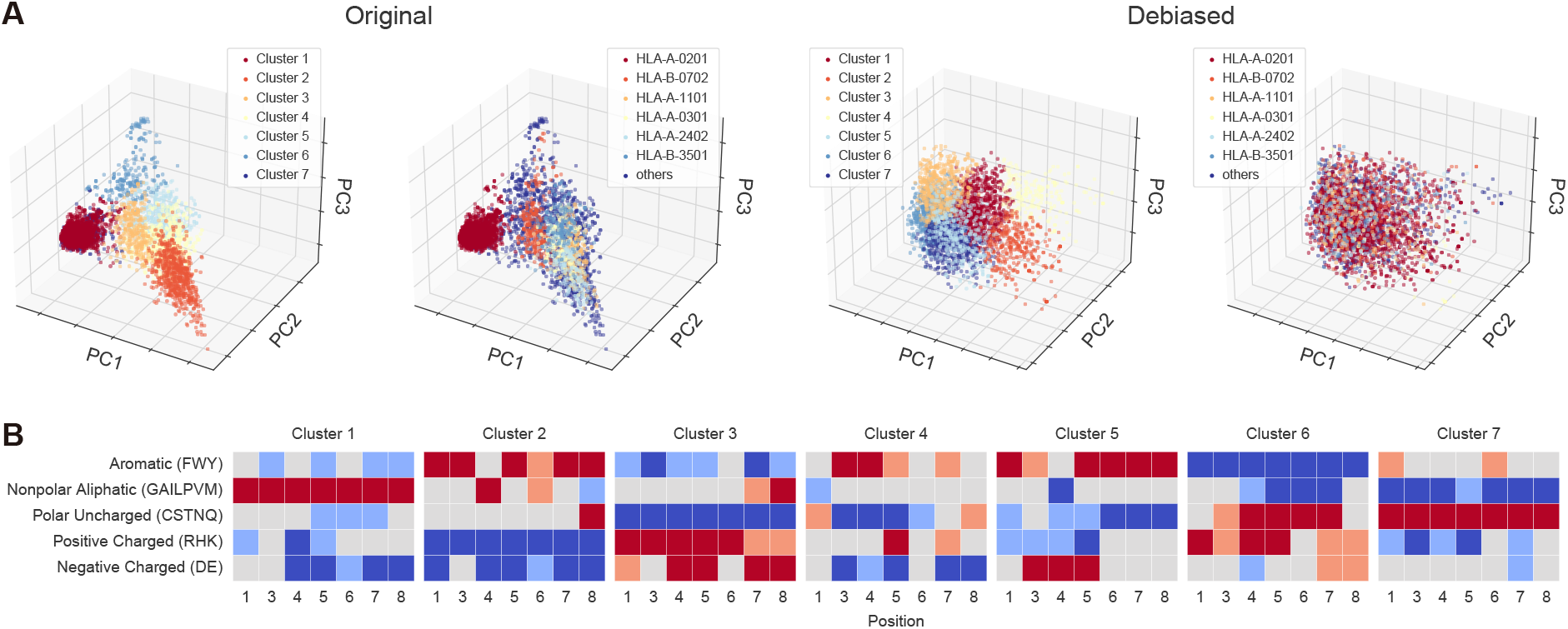
(A) Visualization of immunogenic peptides with model’s embeddings in scatter plots. (B) Statistical analysis related to biophysical properties of amino acids on different positions of different peptide clusters by two-sided binomial test: red - enrichment, *p <* 0.001; light red - enrichment, *p <* 0.05; light blue - depletion; *p <* 0.05; blue - depletion, *p <* 0.001

An important question in immunology is the biophysical patterns associated with immunogenic neoantigens. Previous studies have explored this topic to some extent, suggesting bulk, aromatic, and hydrophobicity are associated with immunogenicity [62, 71]. Twenty amino acids were grouped into five subsets based on their structures and biophysical properties [72]. Then, we conducted a two-sided binomial test to compute the frequency of each amino acid group within each cluster in comparison to the entire population (Fig. 5B). In general, peptides in different clusters demonstrated a strong preference for specific groups of amino acids. For instance, a notable enrichment for nonpolar aliphatic groups appeared in cluster 1, whereas cluster 3 tended to enrich for both positively charged and negatively charged amino acids, along with a depletion of polar uncharged groups. Additionally, aromatic groups and positive groups often shared an adverse trend. These patterns not only are in line with previous studies [62, 71, 66] but also propose an interesting hypothesis that immunogenicity may be influenced by cluster-specific features rather than a universal set of rules applicable to all.

In summary, ImmuBPI_I can efficiently integrate the biophysical properties when encoding peptides. Remarkably, this is achieved with no explicit encoding for their structural and biophysical characteristics as input features to the model.

## 3. Discussion

Neoantigens play a vital role in immunotherapy and to date have demonstrated promising results in clinical trials [30, 31]. Despite various methods that have been developed to predict each individual event in neoantigen immune response, how to identify neoantigens with fewer false positives remains a central challenge [5, 73]. In this paper, we established a unified machine learning framework ImmuBPI that comprised three key steps of the neoantigen immune response process. Through cross-task model interpretability analysis, we have found a significant bias within immunogenicity datasets overlooked in previous research. This has resulted in machine learning models capturing misleading signals regarding the ratio between positive and negative peptides within HLA alleles. We also showed that such a bias would hinder immunogenicity benchmarks from fairly evaluating different models’ performance associated with the learning of immunogenicity. To alleviate this issue, we developed a mutual information-based debiasing solution that successfully disrupted shortcut learning and regained the sensitive immunogenicity prediction for different peptide variants on two case studies. Furthermore, debiased ImmuBPI_I can learn useful representations that encode immunogenicity-related features of the peptides.

In line with previous studies, ImmuBPI demonstrates that binding and presentation prediction can be achieved with high precision. How to prioritize immunogenic epitopes with larger probabilities to stimulate T cell reaction is the key bottleneck in neoantigen identification, which is constrained by insufficient data. Some of the past research focused on strengthening the immunogenicity model by transfer learning from the other two tasks, which could be mainly classified into model-based transfer [28] and feature-based transfer [25, 20, 26]. In this research, we have further discovered and characterized the challenge of data bias which enabled us to rethink the key aspects of training an immunogenicity model. Transfer strategy from previous tasks may deteriorate the learning of immunogenicity by inducing extra bias that further couples with HLA alleles. It also has the potential risk of harming the model’s binding or presentation recognition ability which could be easily learned by a single model.

In light of this, for all the mutated peptide sets generated from sequence alignment between tumors and normal cells, it is recommended that binding and presentation models be utilized first to recall potential peptide binders given specific HLA alleles. Then, immunogenicity models would perform as a ranking module to prioritize possible immunogenic neoantigens. Careful bias control is strongly advised when data-driven methods are employed for immunogenicity tasks to ensure generalization ability at the second stage. By doing so, it is hopeful to realize the full potential of the existing data and models.

In real world applications, especially in biological scenarios, biased data distribution is a common phenomenon. While multiple types of data bias often co-exist in single dataset, determining which bias significantly affects model learning and understanding how it impacts the learning presents a difficult yet meaningful challenge. By utilizing attention mechanism as a powerful model interpreter, we have identified and characterized the challenge of data bias in immunogenicity datasets, particularly the intra-HLA pos/neg sample ratio imbalance. We demonstrated the lack of bias control would lead the model find shortcuts to make predictions, in a model-agnostic way. To the best of our knowledge, while HLA allele frequency imbalance has indeed been widely acknowledged, we are the first to systematically identify and address the bias brought by intra HLA allele imbalance. This is also the first comprehensive analysis highlighting the impact of such biases on immunogenicity prediction models with specific assumption and experimental result.

There are several limitations of this study. The debiasing strategy may not be universally applicable as anchor positions for different HLA alleles could be slightly different. Since HLAs also play a role in TCR interactions, their removal could result in information loss to some extent. It is also important to note that the proposed debiased model was evaluated only on a limited number of retrospective case studies. A larger testing dataset with rigorous bias control is needed to thoroughly benchmark existing methods in the future. Curation for both training datasets and benchmarks with balanced data is of top priority, which may gradually be achieved through the continuous development of high-throughput technologies [74]. As for modeling strategy, incorporating structural information as additional features and utilizing an HLA-adaptive debiasing method with less information loss may lead to improved performance under limited data conditions [75, 76, 77]. In summary, neoantigen-mediated tumor immunotherapy is a promising strategy that requires collaborative efforts from researchers across multiple disciplines.

## 4. Star Methods

### 4.1. Problem formulation

The predictions of the binding, presentation, and immunogenicity are treated in a unified formulation as follow. Let ***x*** represent a sample comprising an HLA-peptide pair from the pair space 𝒳, while *y* ∈ 𝒴 = {0, 1} denotes the corresponding label. It is worth to note that the three distinct processes of interest give rise to three separate dataset distributions *p*_B_, *p*_P_, and *p*_I_ within the space 𝒳 × 𝒴. Hence, our goal was to train three corresponding models *f*_***θ***_ : 𝒳 → [0, 1] to address the classification task for these three processes with cross-entropy loss. Importantly, the binding, presentation, and immunogenicity are prerequisite conditions for subsequent stages. Consequently, the support set *S*_I_ of *p*(***x***, *y* = 1) is contained within *S*_P_ and further within *S*_B_. This proposition will be used in sections of bias analysis.

### 4.2 Attention mechanism of ImmuBPI

As we mentioned in Sec. 2.1, we used an interaction-based transformer encoder to build our framework, ImmuBPI. The centerpiece of this framework is the attention mechanism, which takes the input *X* ∈ ℝ^*l*×*d*^ and an optional attention mask matrix *M* ∈ {0,−∞}^*l*×*l*^ then produces contextual embeddings in 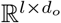 through the adaptive aggregation of information, as equation (1):

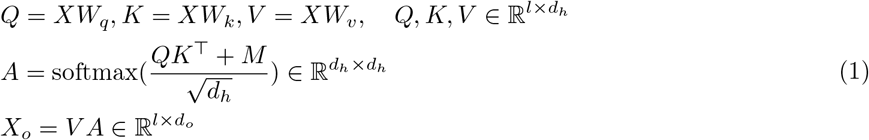

The attention mask is used to block out the *i*th token query the *j*th token in contextual embedding calculation by setting *M*_*i,j*_ = −∞ (vice versa, *M*_*i,j*_ = 0 means allowing the *i*th token to query the *j*th token). In particular, for some of our experiments, we would mask some residue tokens. Mask the *i*th residue token in these experiments just means blocking out all tokens querying the *i*th token by setting *M*_1,*i*_, *M*_2,*i*_, …, *M*_*l,i*_ = −∞.

### 4.3. Mutual information-based debiasing strategy

For generality, we denote *b* as the abstract bias. For instance, in one of our experiments, *b* was the HLA type or the anchor position residues, *y* was the immunogenicity label, and ***x*** was the HLA sequence and peptide sequence pair. In the context of discussing debiasing strategies, it is often observed that the bias feature, *b*, exhibits a strong correlation with the final label *y*, leading to an imbalanced conditional probability *p*(*y* | *b*). However, the distribution of *y* and *b* may significantly shift during real-world testing scenarios. This shift is exemplified by the disparity between *p*_train_(*y, b*) and *p*_test_(*y, b*), rendering any simplistic solutions reliant solely on the influence of *b* ineffective.

In our work, we measured the unbalancedness and detected the bias candidate by mutual information (MI) between *y* and some parts of ***x*** (denoting as *x*^′^), which is empirically useful in different areas [78, 79, 80]. We can consider the marginal-conditional formulation of MI:

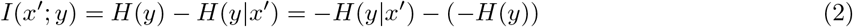

The entropy of a distribution measures its uniformity. Higher entropy values indicate a more uniform and balanced distribution. In other words, greater negative entropy signifies greater distribution imbalance. Therefore, the MI between feature *x*^′^ and label *y* quantifies the increase in imbalance of *y* when transitioning from an unknown *x*^′^ to a given *x*^′^. High MI suggests that feature *x*^′^ is a potential candidate for either being a biased feature or a pivotal feature influencing the label *y*. Since *S*_I_ ⊂ *S*_P_ ⊂ *S*_B_, the label and the high defining features of the previous process could easily be a bias candidate of the process in interest.

After extracting a high MI feature as a bias candidate, we would check the biological causal relationship between it and the ground truth label. Features with weak or counterfactual relationships would be treated as biases and be masked before input in the model as a debiasing strategy.

### 4.4 Training dataset

In this study, we focused on HLA-I alleles, which are presented on the surface of most nucleated human cells. They are responsible for presenting intracellular abnormal peptide epitopes to cytotoxic CD8+ T cells. The training datasets of ImmuBPI used for different tasks are listed in this section. Typically, peptides bound with HLA-I have a length of 8-14 amino acids with a strong preference for 9mers. Peptides over the length of 15 were excluded from training.

#### Binding

We adopted the binding training set from Ref [14], which was originally obtained from Ref [13]. The negative peptide was randomly sampled from IEDB immunopeptidomes based on each binder length and HLA allele, which has proven to have little effect that can be ignored [14]. The training dataset contained 718,332 peptide-HLA pairs. For each binder length and each HLA allele, peptides of negative data are sequence segments that are randomly chosen from the source proteins of IEDB HLA immunopeptidomes.

We kept the same 5-fold split and conducted cross-validation for consistency [14].

#### Presentation

We utilized presentation training sets from Ref [16, 20]. Similar to binding dataset, negative samples were generated by extracting from the UniProt database randomly [81]. Data in mono-allelic samples were reserved. The training dataset contained 4,063,475 peptide-HLA pairs. To ensure a relatively balanced distribution of positive and negative samples, few HLA alleles with only positive or negative samples in the presentation dataset were excluded during training (accounting for only 1.3% of the whole dataset). We conducted a 5-fold cross-validation for consistency [16].

#### Immunogenicity

To make immunogenicity prediction more accurate, we adopted the latest and biggest public training dataset curated by Ref [27]. The training dataset contained 19,933 peptide-HLA pairs. Likewise, only human-related data were reserved. Additionally, HLA alleles that had no corresponding amino acid sequences were either removed or substituted with the HLA allele that had the same 2-digit typing. We conducted a 10-fold cross-validation in consistency with Ref [24]. In section 4.2, to ensure the consistency and fairness necessary for bias exploration and ablation study, we adopted the same dataset curated by Ref [28]. As it is difficult to construct negative samples with high confidence [82], all negative data in immunogenicity dataset are derived from experimental result.

### 4.5. Experimental details

The input HLA was represented by its core sequence, which comprised 34 residues [7], while the peptide was represented by its complete sequence with a maximum length of 15. Each residue was embedded as a learnable embedding. The embeddings of HLA and the embeddings of peptide were added sinusoidal positional embeddings individually. Then, the two embeddings and a learnable [CLS] embedding were concatenated as the input of the backbone model.

To build a unified framework, we used the same hyperparameters and model structures across all three prediction tasks. The models employed in those tasks were pre-LN transformer encoders [83], consisting of 4 layers, 4 attention heads, and 128 hidden states dimensions. The dropout rate between the layers was set at 0.1. The projection head was a ReLU activated 2-layer perceptron, progressively converting hidden state dimensions by 128, 64, 2. We used Adam as the optimizer with a learning rate of *α* = 5 × 10^−5^, and the decayed rate of the first moment and second moment *β*_1_ = 0.9, *β*_2_ = 0.999, respectively. The training process contained 1000 epochs and would be early-stopped when the validation set AUC didn’t increase in 10 epochs. In practice, the convergence was achieved for binding, presentation, and immunogenicity tasks within 310, 80, and 204 epochs respectively. In cases where a model group was trained on cross-validation partitioned datasets, model ensembling was employed by averaging the output probabilities.

### 4.6. Mutation test for model interpretation

For all the 9mer peptide-HLA pairs in the testing benchmark, we mutated the HLA and residue to generate a mutation set. Specifically, every HLA is mutated to all the possible HLA in the respective training set and every peptide residue is mutated to the other 19 amino acids. The abs of score changed upon mutation is recorded and plotted using a violin plot. In principle, a larger value indicates a more significant impact on the model output.

## 4.7. Data and code availability

The data and source code of our work is available at https://github.com/WangLabTHU/ImmuBPI

## Acknowledgments

This work was supported by the National Natural Science Foundation of China (Nos. 62250007 and 62225307), Beijing Municipal Natural Science Foundation (Z230015), and the grant from the Guoqiang Institute, Tsinghua University (2021GQG1023).

## Declaration of generative AI and AI-assisted technologies in the writing process

During the preparation of this work the authors used ChatGPT in order to improve the readability and language. After using this tool, the authors reviewed and edited the content as needed and take full responsibility for the content of the publication.

## Supplementary

**Fig. S1.**
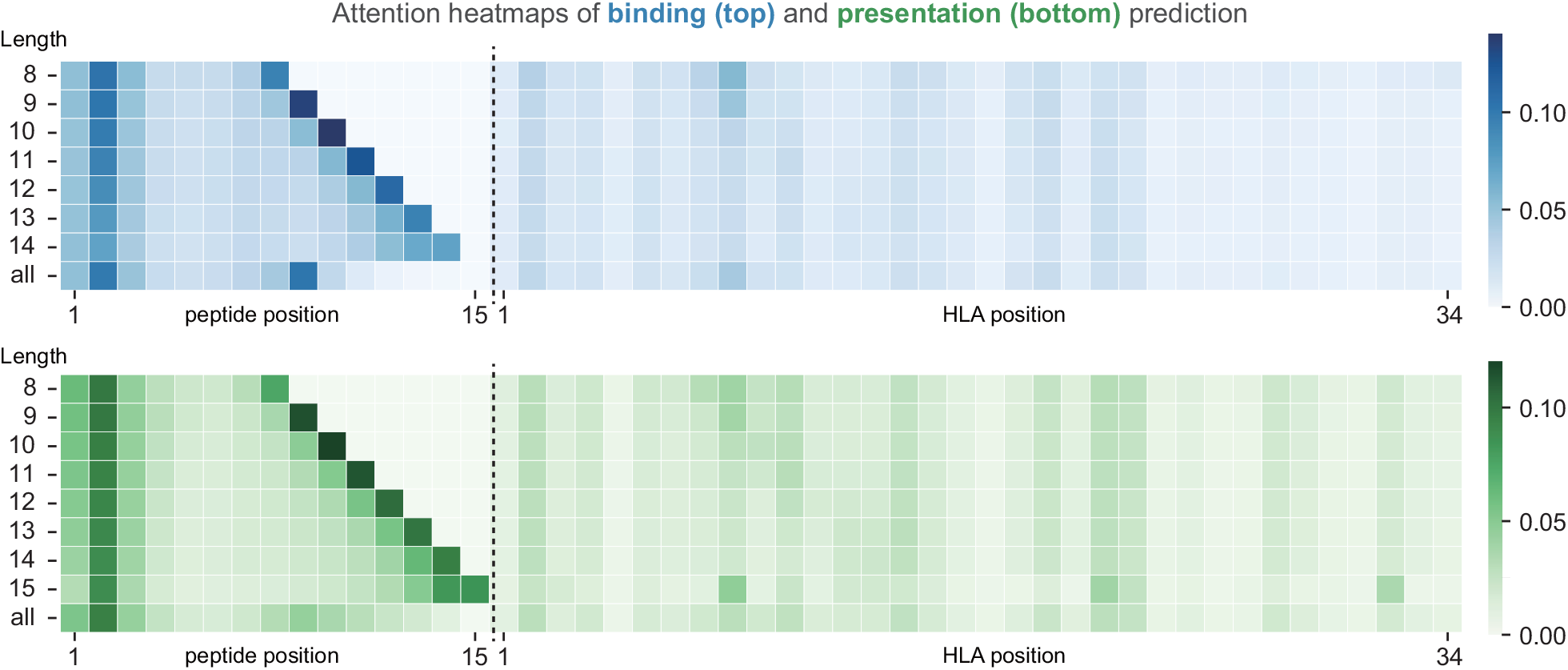
Visualization of attention weights in heatmaps for binding and presentation prediction on peptide groups of different lengths. Calculations were conducted based on the model trained from one of the 5 folds and its respective validation set. Attention weights were averaged between attention layers.

**Fig. S2.**
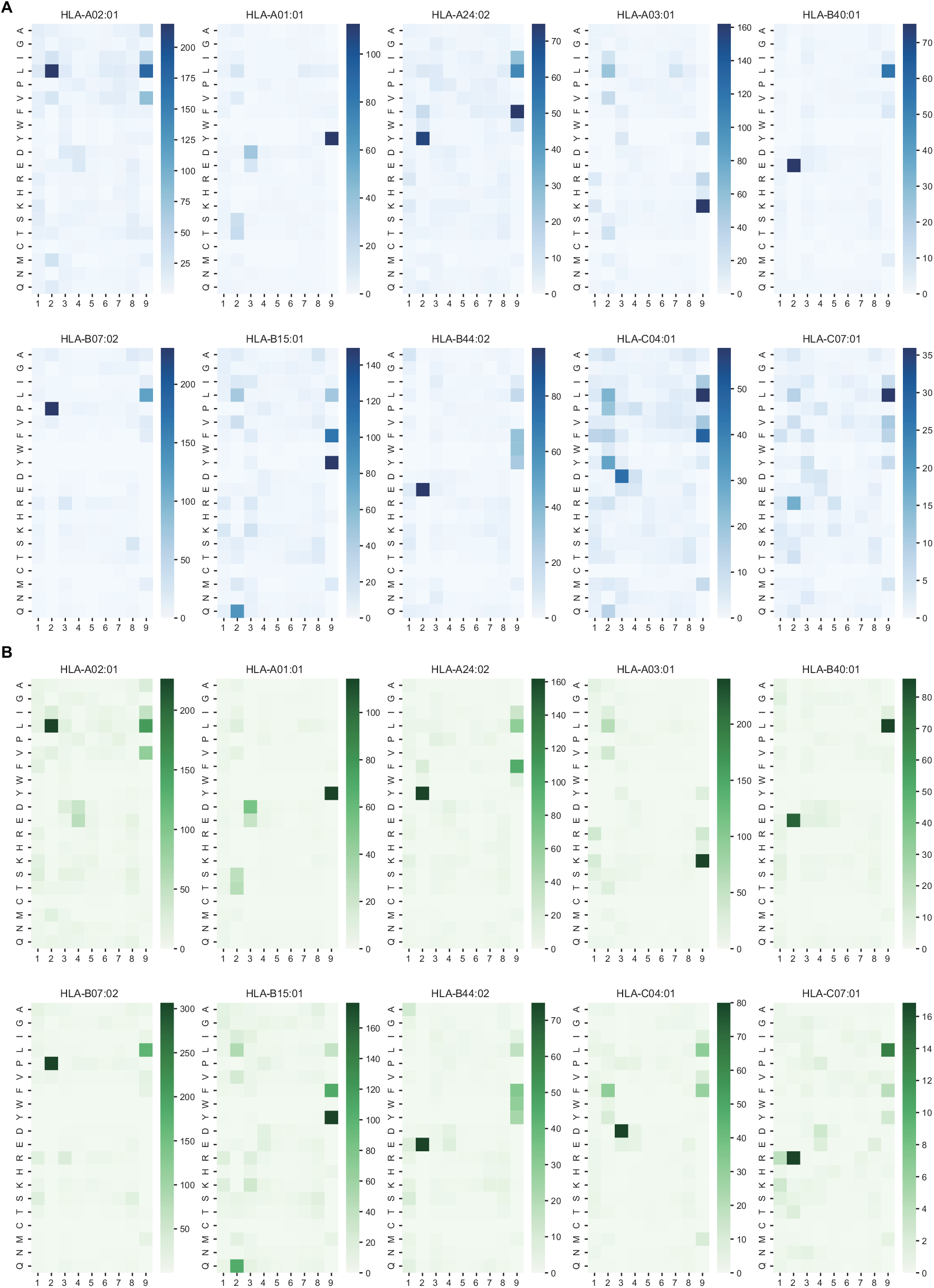
Allele-wise visualization of attention map for binding (A) and presentation (B) prediction. The score was calculated as the sum of the attention weights across 9mer peptides with positive binding or presentation labels. Calculations were conducted based on the model trained from one of the 5 folds and its respective validation set. Attention weights were averaged between attention layers.

**Fig. S3.**
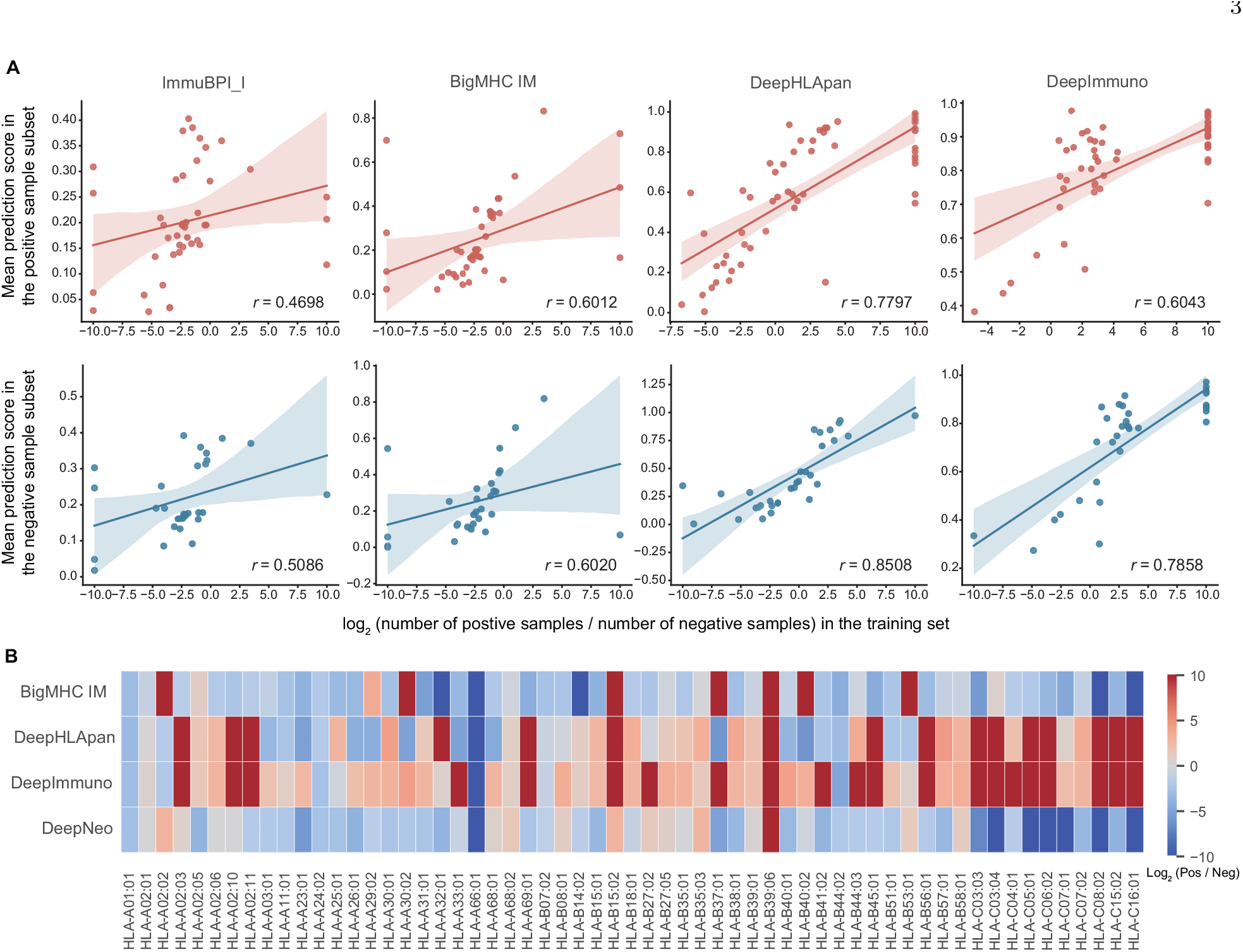
(A) Correlation between model prediction scores in the IEDB infectious disease epitopes benchmark and HLA imbalance degree in the training set. X value was calculated as the log2 ratio between positive and negative peptides corresponding to a given HLA allele in the training set. X value of HLA with only positive peptides was set at 10 while HLA with only negative peptides was set at −10. Y value was calculated as mean prediction scores across all peptide-HLA pairs related to this HLA. HLA with only one sample in the subset was excluded. (B) Log2 ratio between the numbers of positive peptides and negative peptides given HLA allele across different immunogenicity training datasets. The value of HLA with only positive peptides was set at 10 while HLA with only negative peptides was set at −10.

**Fig. S4.**
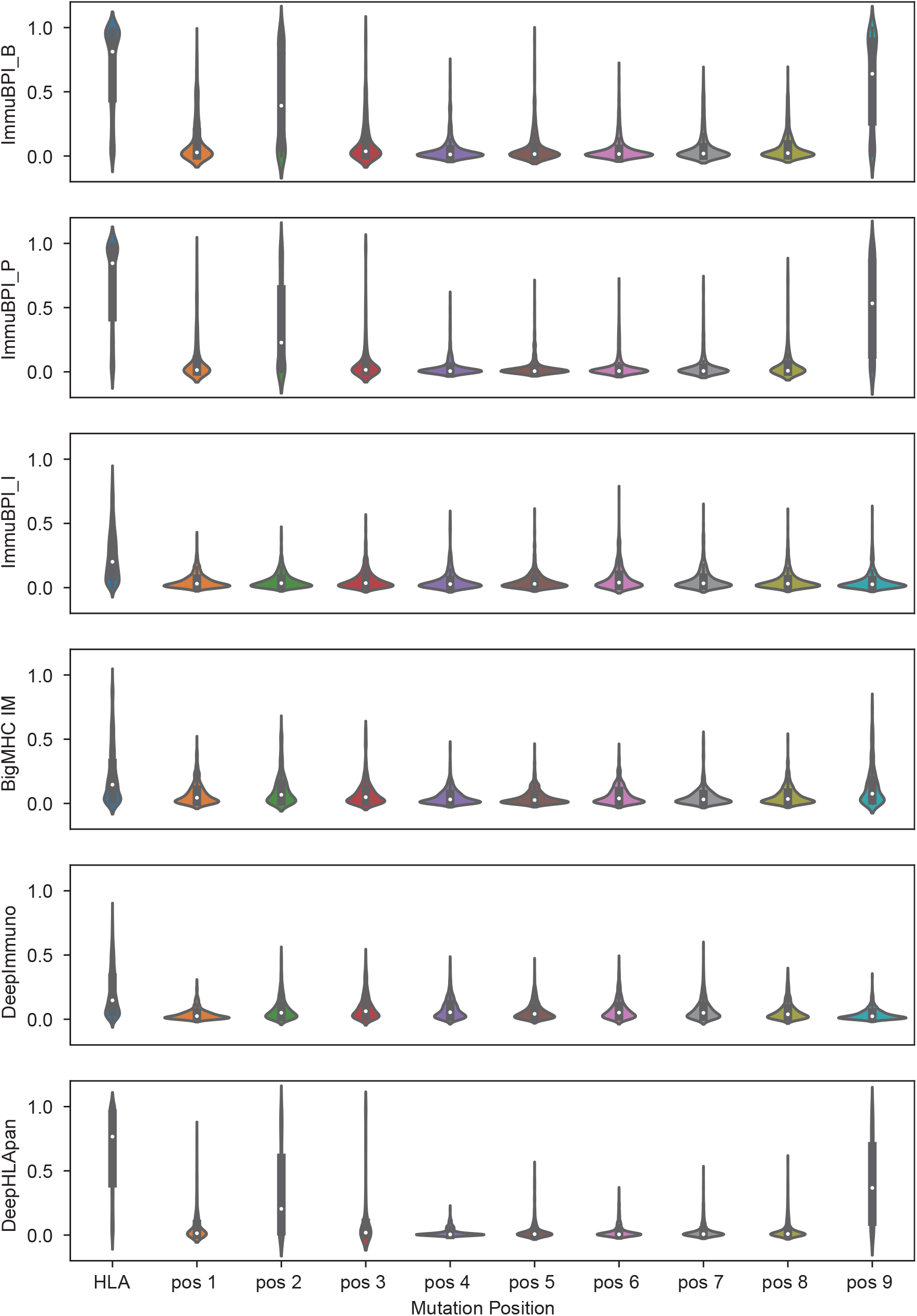
Mutation-based model-agnostic analysis results for four different machine learning models including ImmuBPI_I, BigMHC IM, DeepImmuno and DeepHLAPan.

**Fig. S5.**
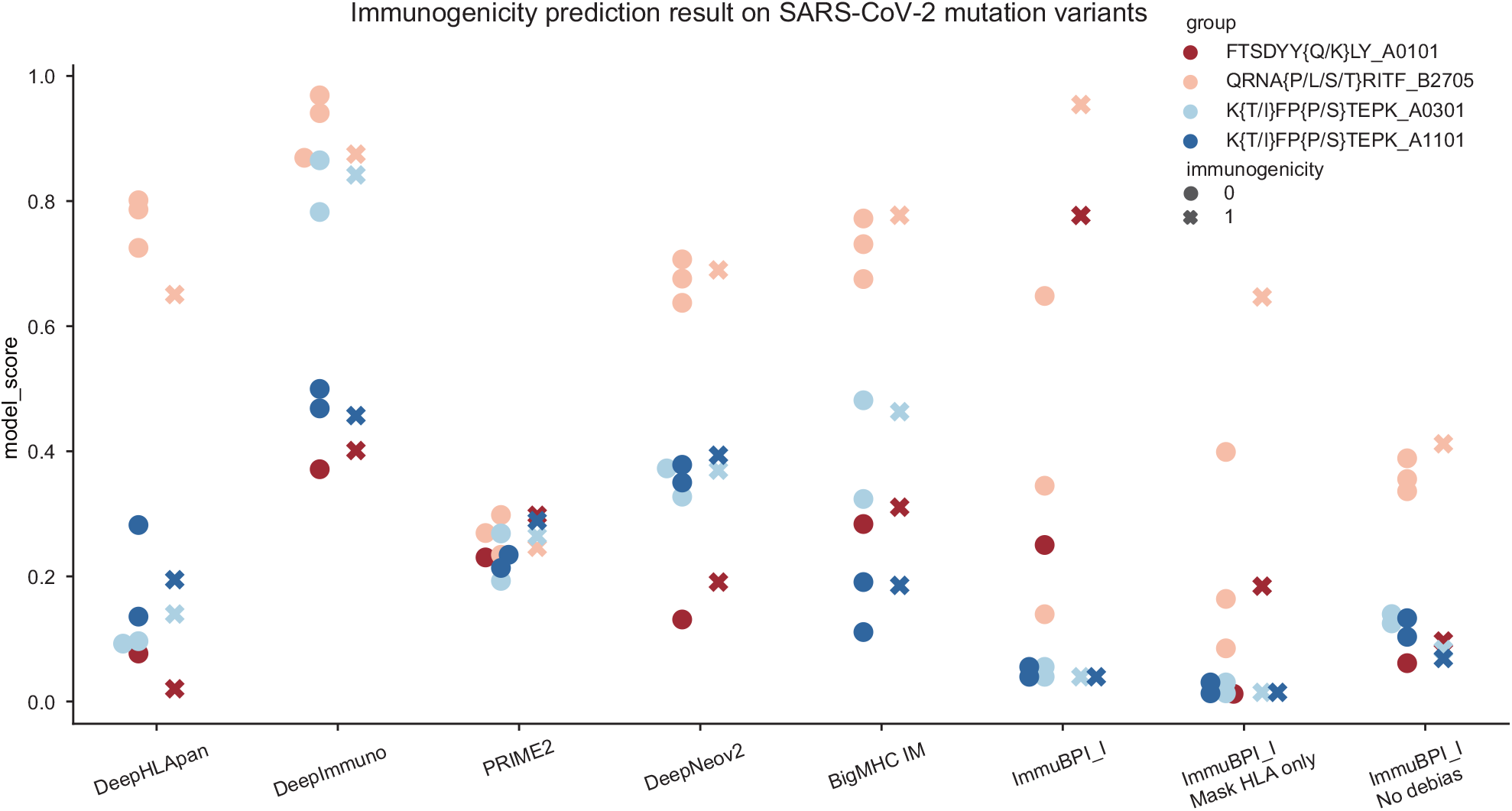
Prediction results of different models on four groups of SARS-CoV-2 single-nucleotide mutation variants.

